# Status of Genome Function Annotation in Model Organisms and Crops

**DOI:** 10.1101/2022.07.03.498619

**Authors:** Bo Xue, Seung Y Rhee

## Abstract

Since entry into genome-enabled biology several decades ago, much progress has been made in determining, describing, and disseminating functions of genes and their products. Yet, this information is still difficult to access for many scientists and for most genomes. To provide easy access and graphical summary to the status of genome function annotation for model organisms and bioenergy and food crop species, we created a web application (https://genomeannotation.rheelab.org) to visualize, search, and download genome annotation data for 28 species. The summary graphics and data tables will be updated semi-annually and snapshots will be archived to provide a historical record of the progress of genome function annotation efforts. Clear and simple visualization of updated gene function annotation, including the extent of what is unknown, will help address the grand challenge of elucidating functions of all genes in organisms.

## Introduction

Rapid advances in DNA sequencing technologies made genome sequences widely available and revealed a plethora of genes encoded within the genomes (O’Leary et al., 2016). The timely invention and wide adoption of the Gene Ontology (GO) system transformed how gene and protein functions are described, quantified, and compared across many organisms (The Gene Ontology Consortium, 2021; Ashburner et al., 2000). A grand challenge in life sciences is to elucidate the functions of all the genes that have been identified through genome sequencing. One of the first steps in elucidating gene function systematically is to know which genes have unknown function. A snapshot of the status of genome function annotation across species is still not readily available for scientists.

Status of genome function annotation is not easily accessible for several reasons. First, genome sequences and their annotations are hosted across multiple databases that use different gene/protein/sequence identifier systems. Although some databases include cross database references and provide tools to map IDs, such as UniProt’s Retrieve/ID mapping and BioMart’s ID conversion (Guberman et al., 2011), these tools are not available for all sequenced genomes. Second, gene function information is not generally annotated using the GO annotation and evidence system in the literature and most databases. Third, genome function annotation databases generally only include annotated genes, and it is often not straightforward to identify unannotated genes. In enrichment analysis, unannotated genes play an important role in reducing ascertainment bias and facilitating discovery of novel processes.

To provide scientists and students an easy way to access and visualize the status of genome function annotations of model species and bioenergy and food crops, we created a web application that displays these data graphically and tabularly and allows searching and downloading annotations using GO term IDs. The website retrieves data from multiple databases, and generates plots that show the percentages of genes with experimental, computational, or no annotations. The snapshots are updated semi-annually, and past snapshots will be archived.

## Results

To represent the status of genome function annotation, we selected three groups of organisms: model organisms, bioenergy model and crop species, and most-annotated plant species (Figure 1). Model organisms are important experimental tools for investigating biological processes and represent key reference points of biological knowledge for other species (Ankeny & Leonelli, 2020; Fields & Johnston, 2005; A. M. Jones et al., 2008). This panel includes: *Arabidopsis thaliana, Caenorhabditis elegans, Danio rerio, Drosophila melanogaster, Mus musculus, Saccharomyces cerevisiae*, and *Schizosaccharomyces pombe* (Fig. 1A). We also included *Homo sapiens*, a species for which many model organisms are studied. Next, we selected bioenergy models and crops, which are important in expanding the renewable energy sector needed to combat the climate crisis and steward a more sustainable environment. Biomass is projected to become the biggest source of primary energy by 2050 (Reid et al., 2020). The bioenergy models and crops we selected include: *Brachypodium distachyon, Chlamydomonas reinhardtii, Glycine max, Miscanthus sinensis, Panicum hallii, Panicum virgatum, Physcomitrium patens, Populus trichocarpa, Sorghum bicolor*, and *Setaria italica* (Fig. 1B). Finally, we included ten additional plant species that have the highest number of GO annotations in UniProt (UniProt Consortium, 2019), which include: *Oryza sativa Japonica Group* (rice), *Gossypium hirsutum* (cotton), *Spinacia oleracea* (spinach), *Zea mays* (corn), *Medicago truncatula, Solanum tuberosum* (potato), *Ricinus communis* (castor bean), *Nicotiana tabacum* (tobacco), *Papaver somniferum* (opium poppy), and *Triticum aestivum* (wheat) (Fig. 1C). These include the world’s most important cereal crops, such as corn, rice, wheat, and vegetable crops such as potato (Food and Agriculture Organization of the United Nations, 2021).

**Figure 1.**
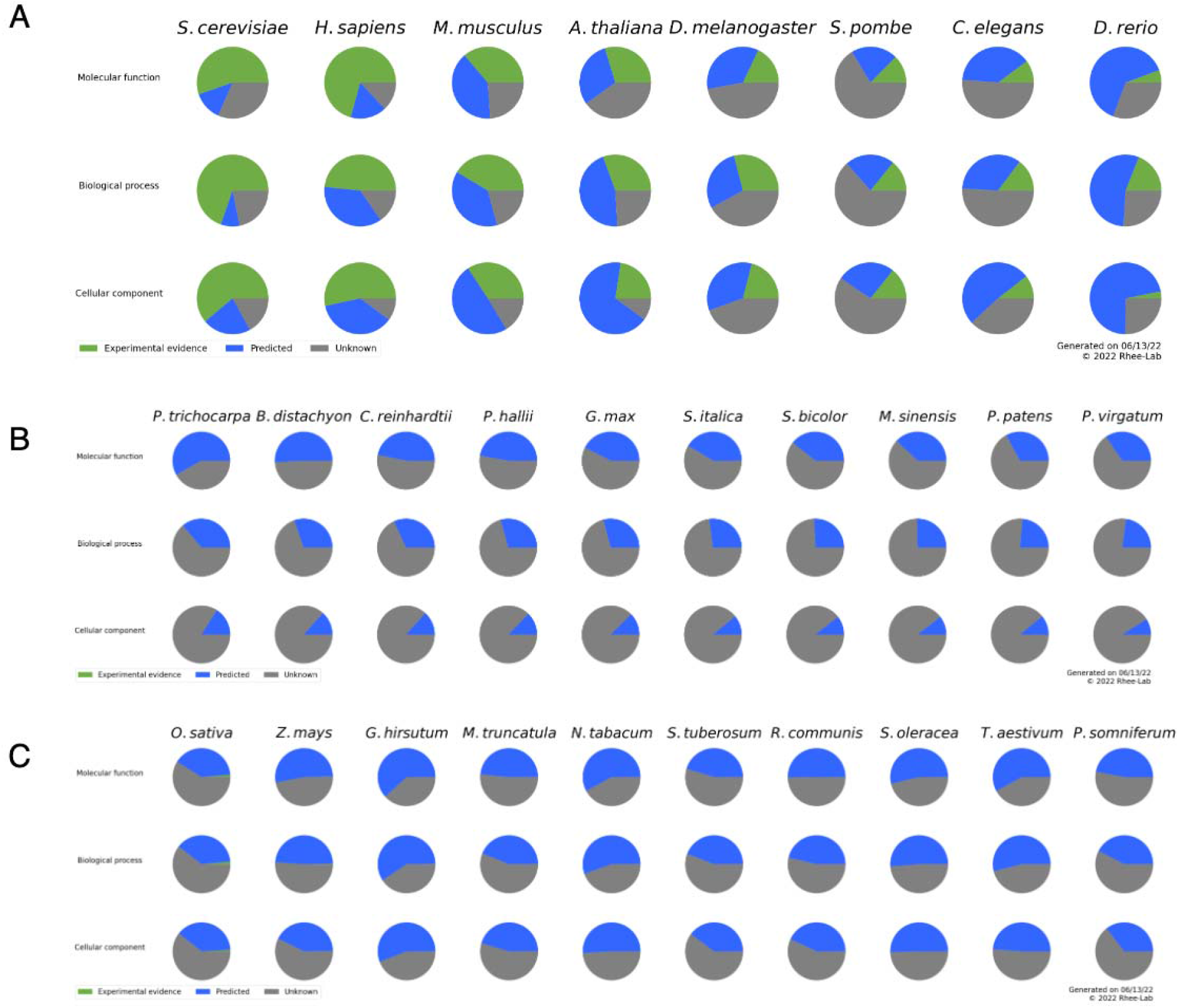
Status of genome function annotations Each pie chart shows the proportion of genes that are annotated to a domain of Gene Ontology (GO): molecular function, biological process, or cellular component. Green indicates genes that have at least one experimentally validated GO annotation; blue indicates genes that are annotated with computationally predicted GO annotations; and gray indicates genes that do not have any GO annotations or have GO annotations with ND (No biological Data available). The species are ordered by the average percentage of genes with experimental evidence (or genes with computationally predicted annotations if experimental evidence is not available) of all three domains. A) selected model organisms; B) bioenergy models and crops; C) other plant species with the highest percentage of genes with experimental evidence in UniProt.

There are several ways of accessing the status of genome function annotation for the 28 species. From the front page, visitors can get a quick summary of the state of the genome function annotation as pie charts for the three groups of species (Figure 1). These pie charts show the percentage of genes that have: 1) annotations with experimental evidence (green); 2) only the annotations that are computationally generated (blue); or 3) no annotations or annotations as being unknown (gray) (Figure 1). Of the eight selected model organisms, *H. sapiens* has the highest percentage of genes with experimental evidence, followed by *S. cerevisiae* and *M. musculus. A. thaliana* has the lowest percentage of “unknown” genes.

Among the model organisms, *C. elegans* is the least known species, with the greatest number of genes unannotated or annotated as having unknown function. Most of the plant species have too few GO annotations based on experimental support to be visible in the pie charts. Visitors can get more detailed information of any of the species by clicking on the species name below the pie charts. Each species page shows additional information about annotation status, including displaying the portion of genes annotated to at least one GO domain (molecular function, cellular component, and biological process (The Gene Ontology Consortium, 2021; Ashburner et al., 2000), as well as a Venn diagram showing the overlap of genes annotated to more than one GO domain (Figure 2). This page also has links to source data and a tabular format of the annotation summary for browsing and downloading. Users can also download lists of annotated genes directly from the summary table or search and download annotations using GO term IDs.

**Figure 2.**
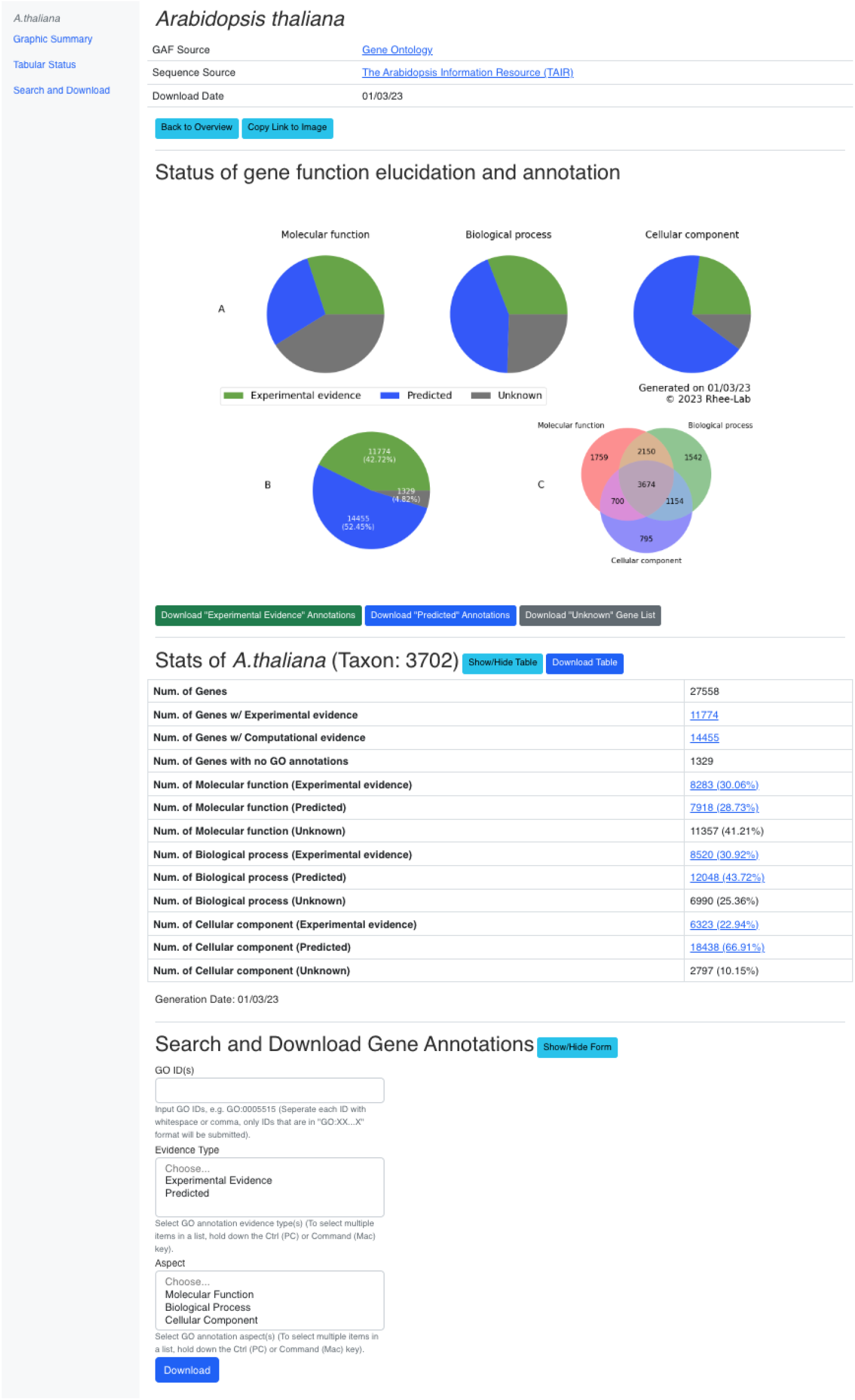
An example species-specific annotation web page shown for Arabidopsis thaliana. It consists of three parts: 1) a table comprising data sources; 2) pie charts showing the proportion of each type of annotations; 3) a table showing the numbers of genes in each category, which users can toggle to either show or hide; and 4) a form to search and download annotations using GO IDs.

## Discussion

Our website provides a convenient way to obtain and visualize the current state of genome function annotation for model organisms and crops for bioenergy, food, and medicine. These charts also serve as a proxy for illustrating how much is known and unknown. These snapshots will be updated on a semi-annual basis, and comparing the charts across time will reflect how biological knowledge changes over time. These snapshots can be useful in many contexts, including research projects, grant proposals, review articles, annual reports, and outreach materials.

In developing our web application, we encountered a few hurdles. First, there was not a single site where all data were available. To obtain GO annotations from the 28 species, we had to visit several databases. An encouraging finding was that all sites that had GO annotations were using the GO Annotation File (GAF) format. Several tools and databases currently provide a single entry point for multiple species GO annotations. AmiGO by the GO consortium (Carbon et al., 2009) and QuickGO from the The Gene Ontology Annotation (GOA) project (Binns et al., 2009) provide search functions for taxon specific GO annotations. However, they require users to set up additional queries to export GAF files of interested species. BioMart from The Ensembl project (Kinsella et al., 2011) is another tool that users can query GO annotations for multiple species using evidence codes. But their output fields do not include publications or references that are linked to the evidence and require users to manually select output fields to conform to the GAF format. Second, our website includes genes that are unannotated, which are often missing in gene function annotations and enrichment analyses (Higgins et al., 2022). Currently, extracting genes that are not annotated requires many steps that differ across species. Including the unannotated genes in a genome into GAF files would facilitate many downstream applications. Third, visualizations of the status of genome annotations do not yet exist, which our website provides using pie charts and tabular summaries.

To our surprise, some plant species with well-maintained, species-specific databases seem to have a low number of experimentally supported GO annotated genes in UniProt. TAIR (Lamesch et al., 2012) and Sol Genomics Network (Fernandez-Pozo et al., 2015) are the only two taxon-specific plant databases that provide GAF files with experimental evidence codes. Most plant genome databases stop at computationally generating GO annotations, and some well-studied species do not appear to have dedicated databases. PLAZA (Van Bel et al., 2022) is a platform that integrates structural and functional annotation for many plant species. They do incorporate experimental evidence from GO (Acids research & 2021, 2021) and GOA (Huntley et al., 2015), but additional annotations are computationally generated using InterProScan or assigned with empirically validated GO annotations from sets of orthologs. More efforts are needed in experimentally validating functional annotations made from computational approaches, characterizing genes of unknown function, and curating experimentally supported function descriptions in the literature into structured annotations such as GO, which will be crucial for accelerating gene function discovery.

The data summarized on this website can be linked to their sources, which can be used for a variety of investigations. Successful examples include exploring why certain proteins remain unannotated (Wood et al., 2019), developing pipelines to infer function without relying on sequence similarity (Bossi et al., 2017), and assessing annotation coverage across bacterial proteomes (Lobb et al., 2020). One of the biggest grand challenges in life sciences today is the limited understanding of what most genes encoded in genomes do. One of the first steps in systemic elucidation of gene function is to know and easily access the set of genes with annotations as well as those without any annotations. Our website provides this functionality for 28 of the most intensely studied species. As our society transitions into biology-enabled manufacturing (Committee on Industrialization of Biology: A Roadmap to Accelerate the Advanced Manufacturing of Chemicals et al., 2015), fundamental knowledge of how genes and their products function at various scales will be crucial, as our society transitions into biology-enabled manufacturing and a new bio-economy era.

## Methods

### Selecting species and data retrieval

For the model organisms, gene function annotations were downloaded as GO Annotation Files (GAF files) from the GO consortium website of the 2022-05-16 release. For *S. Pombe*, data were downloaded from PomBase (Harris et al., 2022) directly. Genes found in a genome were retrieved from the source indicated on the GO annotation download page as General Feature Format (GFF) files. If information of the type of genes are provided in the GFF file, we only included “protein encoding” genes. A detailed description of the files used to generate charts on our website, including data for the other category of species, can be found in Supplemental Table S1.

Apart from *Nicotiana tabacum* and *Papaver somniferum*, all plant species on our website are included in the most recent version of Phytozome V13, but their GO terms are assigned computationally (Goodstein et al., 2012). The Sol Genomics Network (SGN) (Fernandez-Pozo et al., 2015) hosts genome annotations of Solanaceae species, including *Nicotiana tabacum* and *Solanum tuberosum*. An annotation file for *Nicotiana tabacum* is available (Edwards et al., 2017) but they are assigned with computational support coming from InterProScan (P. Jones et al., 2014). SpudDB (Hirsch et al., 2014) provides GO annotation for *Solanum tuberosum* but they are generated with InterProScan and by best hit to the Arabidopsis proteome (TAIR10) (Lamesch et al., 2012). MaizeGDB (Woodhouse et al., 2021) provides GO annotation for *Zea mays* that are assigned with GO annotation tools including Argot2.5, FANN-GO, and PANNZER (Wimalanathan et al., 2018), which are all computational annotations. SpinachBase provides a centralized access to *Spinacia oleracea*, and their GO annotations are generated computationally with Blast2GO (Collins et al., 2019). *Oryza sativa Japonica Group* GO annotations can be found on Rice Genome Annotation Project (Ouyang et al., 2007) and they are assigned with BLASTP searches against Arabidopsis GO-curated proteins (Yuan et al., 2005). Gramene (Tello-Ruiz et al., 2021) hosts genome data for many species but we could not find GO annotations with evidence codes. We were not able to find species-specific databases that provide GO annotations for *Triticum aestivum, Gossypium hirsutum, Medicago truncatula, Papaver somniferum* or *Ricinus communis*.

Genome annotation and gene list for bioenergy models and crops were downloaded from Phytozome version V13. Although some species in this category had GO annotations in the GO consortium database, the sequence identifiers (IDs) for genes could not easily be mapped to Phytozome IDs. To maintain consistency within this category, all annotation files were downloaded from Phytozome. All Phytozome GO annotations are computationally generated (Goodstein et al., 2012). Gene lists were also retrieved from Phytozome V13.

For the last category of plant species, we selected the most annotated plant species from the UniProt GO annotation database (Camon et al., 2004) GAF files hosted on the GO consortium website. We downloaded these species reference proteomes from the UniProt release 2022_02 and retrieved the number of corresponding genes.

Using the evidence codes provided by GAF files, we generated the numbers of genes annotated with GO supported by experimental evidence. If a gene has at least one GO term annotated using any of the following codes: EXP (Inferred from Experiment), IDA (Inferred from Direct Assay), IPI (Inferred from Physical Interaction), IMP (Inferred from Mutant Phenotype), IGI (Inferred from Genetic Interaction), or IEP (Inferred from Expression Pattern), we categorized the gene as having “Experimental Evidence” for function. Genes that have at least one annotated GO term, but no terms have the evidence codes described above, are categorized as “Predicted”. Since Phytozome has only computationally generated GO annotations, all of their genes are categorized as having their functions “Predicted”. By subtracting the annotated genes from the total number of genes, we retrieved the number of genes without any GO annotations. GO annotations with the ND (No biological Data available) are also moved to the “Unknown” category. These numbers were used to generate pie charts to show the proportions of genes in each category for every species.

All files were processed with scripts written in Python (3.10). All pie charts were generated using Python Matplotlib version 3.5.2 and Venn diagrams were generated using Python matplotlib-venn version 0.11.7. The repository of codes can be found at GitHub (https://github.com/bxuecarnegie/AnnotationStats).

## Creating the Website

To create a website for hosting our charts, we used Node.js (Tilkov & Vinoski, 2010) for our server-side environment, which provides the Application Program Interface (API) for the front end to retrieve the plots generated by Python. The front end of the website uses AngularJS (Jain et al., 2014).

## Availability of data and materials

Data used in this study are all publicly available. GO annotation files were downloaded from (http://current.geneontology.org/annotations/index.html 2022-05-16 release, accessed 13 June 2022) and Phytozome (https://data.jgi.doe.gov/refine-download/phytozomeV13, accessed 13 June 2022). Gene data were downloaded from sources indicated on the GO (http://current.geneontology.org/products/pages/downloads.html accessed 13 June 2022), Phytozome, Pombase (http://www.pombase.org/ “pombase-2022-10-01” release) and UniProt (https://www.uniprot.org/ accessed 13 June 2022). Supplemental Table S1 provides detailed information on all species annotation and gene source databases, downloaded versions, and URLs. Scripts used to process the data and generate the graphs are written in Python 3 and are available at GitHub (https://github.com/bxuecarnegie/AnnotationStats).

## Author Contributions

SYR conceived the project, and BX implemented the project. BX and SYR wrote and edited the manuscript.

## Acknowledgments

We thank members of the Rhee Lab for discussions and suggestions on the project and Kristen Yawitz for editing the manuscript. This work was supported, in part, by the U.S. Department of Energy, Office of Science, Office of Biological and Environmental Research, Genomic Science Program grant nos. DE– SC0018277, DE–SC0020366, and DE–SC0021286, and the National Science Foundation grants MCB-2052590, MCB-1916797, and IOS-1546838. This work was done on the ancestral land of the Muwekma Ohlone Tribe, which was and continues to be of great importance to the Ohlone people.

